# Characterization of the RofA regulon in the pandemic M1_global_ and emergent M1_UK_ lineages of *Streptococcus pyogenes*

**DOI:** 10.1101/2023.07.23.550082

**Authors:** Xiangyun Zhi, Ana Vieira, Kristin Huse, Paulo J. Martel, Ludmila Lobkowicz, Ho Kwong Li, Nick Croucher, Ivan Andrew, Laurence Game, Shiranee Sriskandan

**Author notes:** Corresponding author: Professor Shiranee Sriskandan Department of Infectious Disease, Imperial College London, Hammersmith Hospital Campus, Du Cane Road, London W12 0NN, UK.

## Abstract

**Background & Aims:** The standalone regulator RofA is a positive regulator of the pilus locus in *Streptococcus pyogenes*. Found in only certain *emm* genotypes, RofA has been reported to regulate other virulence factors, although its role in the globally dominant *emm*1 *S. pyogenes* is unclear. Given the recent emergence of a new *emm*1 (M1_UK_) toxigenic lineage that is distinguished by three non-synonymous SNPs in *rofA*, we characterized the *rofA* regulon in six *emm*1 strains, that are representative of the two contemporary major *emm*1 lineages (M1_global_ and M1_UK_) using RNAseq analysis, and then determined the specific role of the M1_UK_-specific *rofA* SNPs.

**Results:** Deletion of *rofA* in three M1_global_ strains led to altered expression of 14 genes, including six non-pilus locus genes. In M1_UK_ strains, deletion of *rofA* led to altered expression of 16 genes, including 9 genes that were unique to M1_UK_. Only the pilus locus genes were common to the RofA regulons of both lineages, while transcriptomic changes varied between strains even within the same lineage. Although introduction of the 3 SNPs into *rofA* did not impact gene expression in an M1_global_ strain, reversal of 3 SNPs in an M1_UK_ strain led to an unexpected number of transcriptomic changes that in part recapitulated transcriptomic changes seen when deleting RofA in the same strain. Computational analysis predicted interactions with a key histidine residue in the PRD domain of RofA would differ between M1_UK_ and M1_global_.

**Summary:** RofA is a positive regulator of the pilus locus in all *emm*1 strains but effects on other genes are strain- and lineage-specific, with no clear, common DNA binding motif. The SNPs in *rofA* that characterize M1_UK_ may impact regulation of RofA; whether they alter phosphorylation of the RofA PRD domain requires further investigation.

**Author summary:** RofA belongs to the group of “mga-like” bacterial regulatory proteins that comprise a DNA binding domain as well as a phosphorylation domain (PRD) that is responsive to changes in sugar availability. In certain *emm* genotypes of *Streptococcus pyogenes*, *rofA* sits upstream of the pilus locus, to act as a positive regulator. The recent emergence of a SpeA exotoxin-producing sublineage of *emm*1 *S. pyogenes*, (M1_UK_) has focused attention on the role of RofA; M1_UK_ and its associated sublineages are characterized by 3 non-synonymous SNPs in *rofA,* that include adjacent SNPs in the PRD domain. Here, we determine the impact of *rofA* deletion and the 3 *rofA* SNPs in both the widely disseminated M1_global_ clone and the newly emergent M1_UK_ clone. While production of SpeA undoubtedly contributes to infection pathogenesis, the evolution of M1_UK_ points to a role for metabolic regulatory rewiring in success of this lineage.

## Introduction

Resurgence of scarlet fever and invasive *Streptococcus pyogenes* infection in England has been associated with a sub-lineage (M1_UK_) of the pandemic M1T1 clone, that is distinguished from other prevalent *emm*1 strains by just 27 single nucleotide polymorphisms (SNPs) in the core genome, and increased expression of the phage-encoded superantigen SpeA [1]. Enhanced fitness of M1_UK_ is inferred from its expansion within the population of *S. pyogenes* in England, being detected first in 2010 to becoming dominant by 2015-2016. Among the 27 M1_UK_ lineage defining SNPs, three nonsynonymous mutations were identified in the gene *rofA* encoding the standalone transcriptional regulator RofA, a member of the RALP (RofA-like proteins) transcription regulator family [2]. The SNPs in RofA were also found in the two intermediate *emm*1 sublineages M1_13SNP_ and M1_23SNP_ that accompanied emergence of M1_UK_; M1_13SNP_ was detected as early as 2005, but did not make excess SpeA [3] consistent with a role for a SNP in the leader sequence of ssrA that contributes to the SpeA phenotype, and that is absent in the M1_13SNP_ strains[4].

RofA and its homologue Nra regulate genes of the fibronectin-binding, collagen binding, T-antigen (FCT) region [5]. RofA was first described as a positive transcriptional regulator of the fibronectin binding protein (*prtF*) in *emm6 S. pyogenes*[6]. Reported to bind specific motifs in target promoter DNA, RofA autoregulated its own transcription and appeared to positively regulate the *speB* operon as well as the pilus locus[7]. RofA has however also been reported to negatively regulate a number of other regulators or virulence factors in an *emm* type-specific manner, such as *mga, sagA* and *speA*. RofA negatively regulated *speA* in *emm*6 *S. pyogenes* [2], however, this was not the case for *emm*2 *S. pyogenes* [2, 8].

Despite intense genomic interrogation of the *emm1* lineage and increasing understanding of the genomic wide impact of two component regulators such as covRS (csrRS), little is known about the role of RofA in the *emm*1 lineage. Unlike other genotypes, *prtF* is absent in *emm*1strains, which have a type 2 FCT region, wherein *rofA* is adjacent to the FCT region[9]. The purpose of this study was therefore to characterise the *rofA* regulon in *emm*1 *S. pyogenes* and determine the significance, if any, of the 3 SNPs in RofA detected in the new sub-lineage M1_UK_.

## Results

### 1. Characterization of the RofA regulon in *emm*1 *S. pyogenes*

To systematically characterise the *rofA* regulon in *emm*1 *S. pyogenes,* where the FCT locus is structurally distinct from other genotypes [10] (Figure 1), and to determine the prevalence of strain-specific effects, *rofA* in-frame deletion mutants were constructed in 3 different *emm1* strains H1488, H1489, and H1499 (each of which belong to the globally-disseminated *emm*1 clade designated M1_global_) yielding the isogenic strains M1_H1488rofAKO_, M1_H1489rofAKO_ and M1_H1499rofAKO._

**Figure 1.**
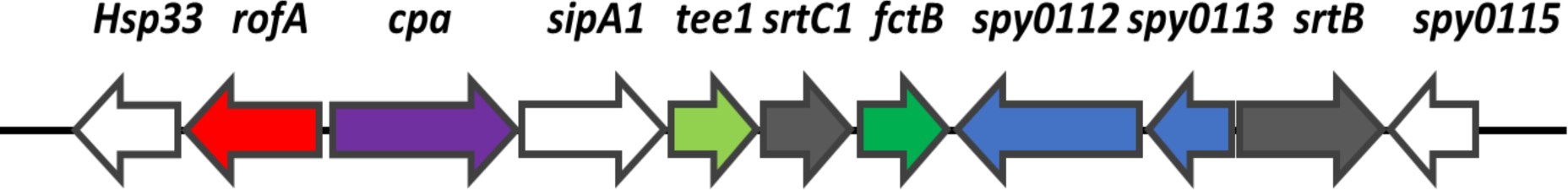
FCT region of *emm*1. Genes are represented by arrows; direction of the arrows indicates direction of transcription. Adapted from Kratovac et al., 2007 [9] and Calfee et al., 2018 [10].

RNAseq analysis of broth-cultured isogenic strains of *S. pyogenes* identified 14 genes that were differentially expressed by at least 1.5-fold in all three isogenic strain pairs. Of these, 11 genes (78.57%) were downregulated in all three of the M1_global_ RofA mutants, consistent with a predominant role for RofA as a positive regulator, while 3 genes (21.43%) were upregulated (Table 1). As expected, transcription of genes associated with collagen binding and the pilus locus (Spy 0107-0114) was markedly downregulated. The effect of RofA mutation on Spy0107 and Spy0109 was confirmed in one panel of isogenic strains by qRT-PCR; mRNA transcripts were significantly reduced in the absence of RofA, and this was reversed by complementation (Figure S1A and B). Spy1081 (PTS system, cellobiose-specific IIA component), Spy1281 (two-component response regulator), and Spy1453 (phage protein) were also downregulated in all three *rofA* mutants. Genes that were upregulated in all *rofA* mutants included Spy0212 (N-acetylmannosamine-6-phosphate 2-epimerase), Spy0213 (N-acetylneuraminate-binding protein) and *scpA*, the gene encoding C5a peptidase. RNAseq failed to show any impact of *rofA* deletion on *speA* transcription. To more rigorously establish that there was no effect of RofA on SpeA, we quantified the transcription of *speA* in M1_H1488rofAKO_ using qRT-PCR but observed no difference in comparison with the parent strain H1488 (Figure S2A).

**Table 1.** Genes differentially regulated in all three M1_globalrofAKO_ strains compared with isogenic parent M1_global_ strains.

| Gene ID | Gene Name | Description | H1488 |  | H1489 |  | H1499 |  |
| --- | --- | --- | --- | --- | --- | --- | --- | --- |
|  |  |  | log2fold-change | padj | log2fold-change | padj | log2fold-change | padj |
| Spy0106 | rofA | transcriptional regulator | -11.241 | <0.001 | -13.902 | <0.001 | -13.938 | <0.001 |
| Spy0107 | NA | - | -3.388 | <0.001 | -3.747 | <0.001 | -3.816 | <0.001 |
| Spy0108 | NA | signal peptidase I | -3.348 | <0.001 | -3.510 | <0.001 | -3.952 | <0.001 |
| Spy0109 | NA | - | -3.288 | <0.001 | -3.514 | <0.001 | -3.983 | <0.001 |
| Spy0110 | eft-LSL | putative exported protein | -3.478 | <0.001 | -3.399 | <0.001 | -4.363 | <0.001 |
| Spy0111 | NA | hypothetical protein | -3.406 | <0.001 | -3.596 | <0.001 | -4.044 | <0.001 |
| Spy0113 | NA | transposase | -2.674 | <0.001 | -2.463 | 0.001 | -1.153 | <0.001 |
| Spy0114 | NA | sortase | -2.248 | <0.001 | -1.710 | <0.001 | -0.612 | 0.002 |
| Spy0212 | NA | N-acetylmannosamine-6-phosphate 2-epimerase | 0.723 | 0.002 | 1.855 | 0.001 | 1.417 | <0.001 |
| Spy0213 | NA | N-acetylneuraminate-binding protein | 0.812 | 0.002 | 1.936 | 0.026 | 1.506 | <0.001 |
| Spy1081 | NA | PTS system, cellobiose-specific IIA component | -0.755 | 0.019 | -0.952 | 0.010 | -1.270 | <0.001 |
| Spy1281 | NA | two-component response regulator | -0.601 | 0.010 | -0.579 | 0.031 | -0.601 | <0.001 |
| Spy1453 | NA | phage protein | -0.694 | 0.005 | -0.676 | 0.033 | -0.804 | <0.001 |
| Spy1715 | scpA | C5A peptidase precursor | 0.589 | 0.003 | 2.323 | <0.001 | 0.510 | 0.010 |

The clear role of RofA in regulation of the pilus locus in the three different M1_global_ strains concealed quite marked inter-individual differences between strains in the genes regulated by *rofA* (Tables S2-4). Indeed, the number of genes downregulated by *rofA* mutation ranged from 60-146, while the number of genes upregulated ranged from 95-264 when considering individual isogenic strain pairs. In some cases, genes that were downregulated in one strain, were upregulated in another. Notably only one *rofA* mutant showed upregulation of *speA* transcription, a derivative of H1499, which showed a 1.99-fold increase in *speA* transcription.

There were however some notable similarities in the *rofA* regulon between pairs of M1_global_ strains (Tables S5, S6, S7). For example, when considering M1_global_ strains H1489 and H1499, (Table S6) streptolysin O and *nga* were upregulated in the absence of functional *rofA*, as were the two phage-encoded DNAses, Spd3 and sdaD2. However, the magnitude of effect was much greater in H1489; there was a 5-6 fold increase in transcription of phage encoded DNAses in the absence of *rofA,* and an 8-fold increase in *slo* transcription.

When considering M1_global_ strains H1488 and H1499, we found that *rofA* mutation resulted in downregulation of the entire operon Spy1732-1736 comprising *speB* and genes concerned with its processing and export, as well as the adjacent gene encoding DNAseB. This amounted to a 9-fold reduction in *speB* in H1488 and a 3.2-fold reduction in *speB* in H1499. This effect of RofA on *speB* was not observed at all in the H1489 *rofA* mutant; indeed, in this strain *rofA* mutation led to a >2-fold increase in Spy1738 *spd* (DNaseB). This strain also demonstrated a convincing 3-4-fold upregulation of the entire SLS operon when *rofA* was mutated, alongside a 3-5-fold increase in genes that comprise the *mga* regulon including *scpA*, *sic* and *emm*. There was no obvious reason for the divergence between strains in the components of the *rofA* regulon, in particular there were no obvious variants in known regulatory genes.

### 2. Impact of 3 SNP RofA mutation in M1_global_ strains

There are 3 non-synonymous mutations in *rofA* that are characteristic of all of the recognized intermediate *S. pyogenes emm*1 sublineages that preceded or accompanied emergence of M1_UK_ [3]. To determine if these SNPs (M318I, F319V, D491N), two of which result in adjacent amino acid changes, have a measurable impact on *S. pyogenes* gene regulation, we introduced the same 3 SNPs into M1_global_ strain H1488 to generate GAS-M1_H1488rofA3SNPs_ (strain designation H1666). RNAseq was then used to compare the transcriptome of the transformant H1666 and the parent strain when cultured in broth in identical conditions to those above. Intriguingly, no genes were found to be differentially regulated using a threshold of log_2_ 1. Three genes were detected as DE with a log_2_ fold change of 0.5 (Spy0501; Spy1678; *smeZ*) but these did not overlap with transcripts affected by *rofA* deletion. Furthermore, there was no impact on pilus locus genes when qRTPCR was used (Figure S1A and B).

Growth of the *rofA* mutant strains, and those with the 3SNPs introduced was evaluated in chemically defined medium but differences identified between the *rofA* mutants and parent strains were minimal (Figure S3 A and B). We considered the possibility that deletion of *rofA* might impact growth of *S. pyogenes* in other more relevant media, however growth and survival of the three RofA mutant strains in whole human blood did not differ from the isogenic parent strains (Figure S4).

As the main reservoir for *S. pyogenes* is the nasopharynx, we compared the ability of the panel of three isogenic *rofA emm*1 strains to cause experimental nasopharyngeal infection using an established mouse model. Detection of a difference in longevity of carriage of *emm*1 isogenic strains was challenging due to severity impacting group size over time despite the low inoculum volume. There was an apparent trend for mice infected with *S. pyogenes* carrying the *rofA* 3SNP (GAS-M1_H1488rofA3SNPs_) to cause more intense infection lasting up to 7 days (Figure S5 A, B and C).

### 3. Characterization of the *rofA* regulon in emergent *S. pyogenes* lineage M1_UK_

We considered the possibility that the function of *rofA* may differ between *emm*1 lineages, and therefore undertook *rofA* gene deletion in a panel of three M1_UK_ strains (H1496, H1491 and H1490) to yield M1_H1496rofAKO_, M1_H1491rofAKO_ and M1_H1490rofAKO_. RNAseq comparison of the 3 pairs of isogenic strains cultured in broth revealed that 16 genes were differentially regulated in all 3 pairs of M1_UKrofAKO_ relative to the M1_UK_ parent strains (Table 2). Of these, 15 genes (93.8%) were positively regulated by RofA, including the FCT locus genes, *bglA*2 operon (cellobiose PTS transporter operon), Spy1732 (protein export protein prsA precursor), and Spy1736 (hypothetical protein). A single gene, *glnQ*.2, was negatively regulated. Genes that were specifically DE in all three M1_UK_ strains (i.e. were not observed following *rofA* deletion in M1_global_ strains) included *epf*, *bglA2* operon, the *prsA* precursor, and a hypothetical protein Spy1736.

**Table 2.** Genes differentially regulated in all three M1_UKrofAKO_ strains compared with parent M1_uk_ strains.

| Gene ID† | Gene Name | Description | H1496 |  | H1491 |  | H1490 |  |
| --- | --- | --- | --- | --- | --- | --- | --- | --- |
|  |  |  | log2fold-change† | padj | log2fold-change† | padj | log2fold-change† | padj |
| Spy0106 | rofA | transcriptional regulator | -9.693 | <0.001 | -12.762 | <0.001 | -14.182 | <0.001 |
| Spy0107 | NA | - | -3.391 | <0.001 | -4.834 | <0.001 | -4.065 | <0.001 |
| Spy0108 | NA | signal peptidase I | -3.220 | <0.001 | -5.301 | <0.001 | -4.389 | <0.001 |
| Spy0109 | NA | - | -3.486 | <0.001 | -5.181 | <0.001 | -4.517 | <0.001 |
| Spy0110 | eft-LSL | putative exported protein | -3.305 | 0.001 | -5.348 | <0.001 | -4.144 | <0.001 |
| Spy0111 | NA | hypothetical protein | -3.622 | <0.001 | -5.206 | <0.001 | -3.966 | <0.001 |
| Spy0113 | NA | transposase | -2.405 | <0.001 | -1.771 | 0.002 | -2.179 | 0.002 |
| **Spy0561** | **epf** | **putative extracellular matrix binding protein** | **-0.851** | **0.002** | **-0.586** | **0.008** | **-1.038** | **0.010** |
| **Spy1077** | **glnQ<sub>2</sub>** | **glutamine transport ATP-binding protein** | **0.796** | **0.022** | **0.771** | **0.001** | **1.334** | **0.005** |
| **Spy1079** | **NA** | **PTS system 2C cellobiose-specific IIC component** | **-1.572** | **<0.001** | **-0.789** | **0.001** | **-0.994** | **0.004** |
| **Spy1080** | **NA** | **hypothetical protein** | **-1.275** | **0.002** | **-0.712** | **0.010** | **-1.108** | **0.006** |
| **Spy1083** | **NA** | **transcription anti-terminator 2C BglG family** | **-1.116** | **0.002** | **-0.589** | **0.002** | **-0.943** | **0.003** |
| **Spy1084** | **NA** | **outer surface protein** | **-1.099** | **<0.001** | **-0.821** | **0.003** | **-1.030** | **0.002** |
| **Spy1085** | **bglA<sub>2</sub>** | **Beta-glucosidase** | **-1.218** | **0.014** | **-0.758** | **0.005** | **-0.996** | **0.004** |
| **Spy1732** | **NA** | **protein export protein prsA precursor** | **-1.518** | **0.025** | **-0.722** | **0.002** | **-0.789** | **0.003** |
| **Spy1736** | **NA** | **hypothetical protein** | **-2.023** | **0.004** | **-1.420** | **0.001** | **-0.688** | **0.028** |

† differentially expressed genes that are specific to M1<sub>UK</sub> are in bold.

Considering the individual pairs of isogenic strains, *rofA* deletion led to downregulation of between 73-124 genes in the three different M1_UK_ strains, and upregulation of 46-174 genes (Table S8, S9, S10). Although there was much interindividual strain variation among the M1_UK_ strains tested, there were again additional notable similarities in the *rofA* regulon between pairs of M1_UK_ strains (Tables S11, S12, S13). Two M1_UK_ strains (H1496 and H1491, Table S11) showed 2-5 fold upregulation of streptolysin O expression and downregulation of SPEB and adjacent genes following *rofA* disruption, similar to results observed among M1_global_ strains. RofA inactivation in the two M1_UK_ strains H1496 and H1490 (Table S12) resulted in downregulation of two operons: the V-type sodium ATP synthase *ntp* operon, and the citrate lyase *cit* operon. Neither operon was downregulated among M1_global_ strains. Interestingly, RofA inactivation in M1_UK_ strain H1491 had an opposing effect on both the *ntp* operon and citrate lyase operon, both of which were upregulated in response to *rofA* mutation, along with upregulation of the *pur* operon (Spy0022-0034), again illustrating quite marked inter-individual strain variation in the impact of *rofA* mutation even among strains that are seemingly phylogenetically related.

There was a substantial overlap in 7 genes that were positively regulated by RofA in both M1_global_ and M1_UK_ lineages, all of which were part of the FCT locus operon. Gene expression differences were confirmed by qRT-PCR (Figure S1). We conclude that these genes therefore represent the core RofA regulon in *emm*1 *S. pyogenes* (Table 3). Similar to earlier findings, there was no evidence that *speA* was regulated by RofA in the M1_UK_ lineage, as confirmed by qRT-PCR (Figure S2 B). Taken together the results pointed to a diversified RofA regulon in M1_UK_ strains. Analysis of promoter regions of genes regulated by RofA in at least 2 of the 3 strains of each lineage (M1_UK_ and M1_global_) was performed and no common motif was identified, even when genes up- and downregulated were considered separately (Figure S6).

**Table 3.**
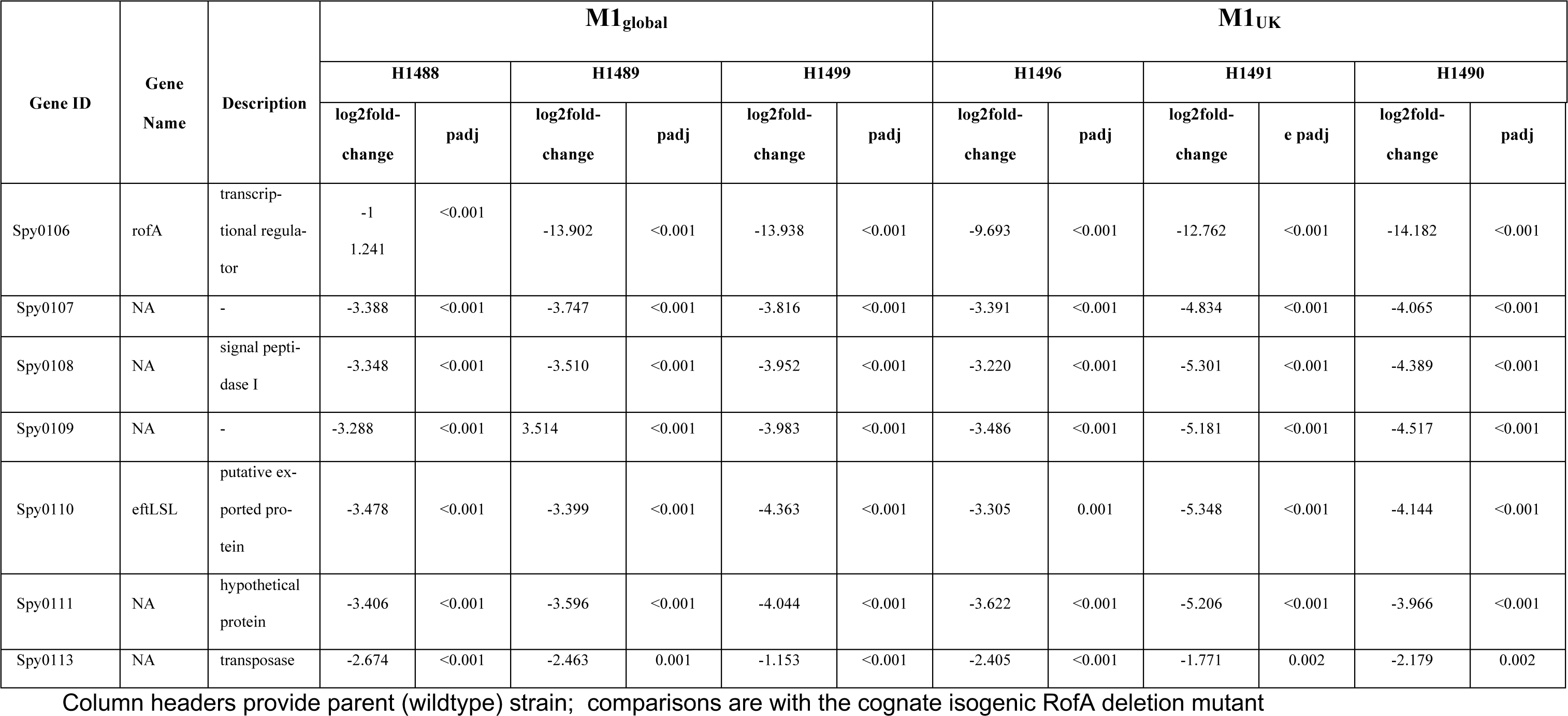
RofA core regulon. Genes differentially expressed in both M1_globalrofAKO_ and M1_UKrofAKO_ strains compared with parent M1_global_ and M1_UK_ strains.

### 4. Reversal of the RofA 3SNPs in M1_UK_

As the RofA regulon in M1_UK_ appeared distinct from M1_global_, it seemed possible that the 3SNPs present in *rofA* in the M1_UK_ lineage may be significant in this strain background, noting that the strain background is characterized by a number of additional SNPs that may alter strain physiology. We therefore evaluated the impact of ‘reversing’ the *rofA* 3SNPs in M1_UK_ strain H1496, to result in strain M1_UKrofA3SNPsFixed_ (strain designation H1665). In contrast to our findings in M1_global_, where introduction of the 3SNPs made little impact on the transcriptome, we found that 91 genes were differentially regulated in the M1_UKrofA3SNPsFixed_. These genes included bacteriocin, *epuA*, 8 phage genes, and several hypothetical genes (Table S14). There was surprising overlap between the transcriptome of the RofA deletion mutant and the mutant with reversal of the 3SNPs in the same strain background (Figure 2), suggesting that the function of RofA, at least in this strain, was impaired by reversal of the 3SNPs. Two genes (Spy1142-1143) were downregulated in both M1_UKrofAKO_ and M1_UKrofA3SNPs_. Over 40% (27 /66) (Table 4) of the genes upregulated in the same M1_UK_ *rofA* mutant were also upregulated when the 3SNPs in *rofA* were ‘fixed’; these included three cytosolic proteins, two phage proteins, three LSU ribosomal proteins, *epuA*, as well as iojap protein family protein and *nrdG*. None of these genes were part of the core RofA regulon. The findings attribute a role for the 3SNPs in RofA repressor activity in M1_UK_ strains.

**Figure 2.**
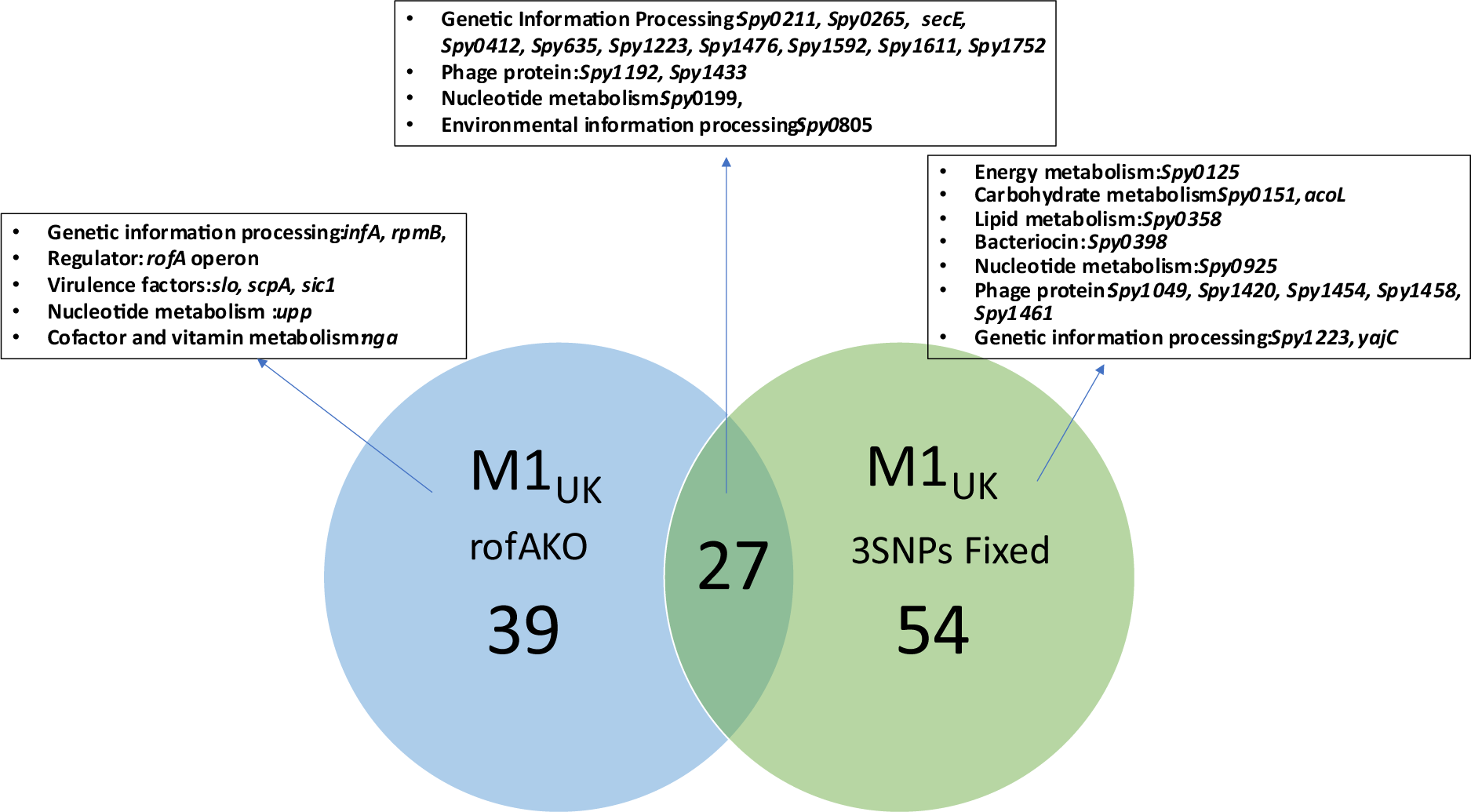
Venn diagram enumerating upregulated genes following disruption of rofA (blue) and replacement of 3SNPs (green) in isogenic derivatives of M1_UK_ strain H1496. Left circle, M1UK_rofAKO_, right circle: M1_UK3SNPsfixed_. Number of overlapping genes shown in centre. Representative genes assigned to each category are indicated in boxes.

**Table 4.** Genes differentially expressed in both M1_UKrofA3SNPsFixed_ and M1_UKrofAKO_ compared with parent M1_UK_ strain.

| Gene ID | Gene Name | Description | M1<sub>UKrofA3SNPsFixed</sub> |  | M1<sub>UKrofAKO</sub> |  |
| --- | --- | --- | --- | --- | --- | --- |
|  |  |  | log2fold-change | padj | log2fold-change | padj |
| Spy0073 | NA | hypothetical protein | 1.334 | 0.003 | 1.305 | 0.007 |
| Spy0199 | NA | deoxyuridine 5'-triphosphate nucleotidohydrolase | 0.752 | 0.003 | 0.828 | 0.045 |
| Spy0211 | rpmH | LSU ribosomal protein L34P | 1.483 | 0.015 | 1.669 | 0.003 |
| Spy0265 | NA | iojap protein family | 1.423 | 0.003 | 1.510 | <0.001 |
| Spy0269 | NA | putative cytosolic protein | 1.213 | 0.003 | 1.260 | 0.001 |
| Spy0399 | NA | putative membrane associated protein | 3.000 | 0.005 | 2.556 | 0.003 |
| Spy0412 | NA | LSU ribosomal protein L33P | 1.793 | 0.002 | 1.671 | 0.008 |
| Spy0493 | NA | putative cytosolic protein | 1.494 | 0.002 | 1.154 | 0.011 |
| Spy0585 | epuA | epuA protein | 1.392 | 0.003 | 1.176 | 0.002 |
| Spy0635 | rpmA | LSU ribosomal protein L27P | 0.710 | 0.020 | 0.659 | 0.037 |
| Spy0805 | srtK | nisin biosynthesis sensor protein | 0.655 | 0.011 | 0.680 | 0.017 |
| Spy0985 | NA | phnA protein | 0.947 | 0.006 | 0.894 | 0.012 |
| Spy1087 | NA | putative cytosolic protein; | 1.889 | 0.005 | 2.208 | <0.001 |
| Spy1089 | NA | hypothetical protein | 2.783 | 0.001 | 2.150 | 0.005 |
| Spy1192 | rsuA | phage protein | 2.609 | 0.001 | 1.689 | 0.015 |
| Spy1223 | NA | DNA-binding protein HU | 1.011 | 0.004 | 0.991 | 0.015 |
| Spy1239 | NA | putative cytosolic protein | 1.062 | 0.001 | 1.447 | 0.014 |
| Spy1295 | NA | putative cytosolic protein | 0.814 | 0.016 | 1.084 | 0.015 |
| Spy1433 | NA | phage protein | 1.361 | 0.004 | 1.286 | 0.007 |
| Spy1476 | NA | ATP/GTP hydrolase | 0.882 | 0.011 | 1.165 | 0.001 |
| Spy1504 | NA | putative membrane spanning protein | 1.428 | 0.002 | 1.175 | 0.010 |
| Spy1592 | NA | ribosomal-protein-alanine acetyltransferase | 0.612 | 0.032 | 0.691 | 0.015 |
| Spy1611 | rpoE | DNA-directed RNA polymerase delta chain | 0.646 | 0.023 | 0.947 | 0.001 |
| Spy1650 | NA | degV family protein | 1.274 | 0.003 | 0.886 | 0.046 |
| Spy1752 | NA | LSU ribosomal protein L33P | 1.849 | 0.001 | 1.287 | 0.032 |
| Spy1789 | nrdG | anaerobic ribonucleoside-triphosphate reductase activating protein | 0.643 | 0.027 | 0.738 | 0.009 |
| Spy1792 | NA | hypothetical protein | 1.293 | 0.003 | 1.368 | 0.008 |

We considered the possibility that the activity of RofA may impact M1_UK_ *S. pyogenes* growth but did not detect a significant difference when comparing growth of a panel of isogenic strains in CDM (Figure S3C and D). We again compared the survival of three M1_UK_ parent strains, with the three isogenic M1_UK_ *rofA* mutant derivatives in whole human blood, however found no difference between the isogenic strains (Figure S4). Finally, we were unable to detect a difference when comparing the strains in a murine model of nasopharyngeal carriage albeit that again, group sizes were affected by infection severity (Figure S7A, B and C).

### 5. Bioinformatic analysis of RofA and implications of the 3SNPs

To predict if the location of the 3 non-synonymous SNPs would have consequences for RofA regulatory function, bioinformatic analysis of the RofA amino acid sequence was carried out. Despite the low genetic similarity (and 19-20% amino-acid sequence identity), domain organization of RofA was remarkably similar to AtxA, a virulence regulator from *Bacillus anthracis* (4R6I); MgaSpn, a Mga regulator from *Streptococcus pneumoniae* (5WAY); and putative Mga family transcriptional regulator from *Enterococcus faecalis* (3SQN) as shown in Figure 3. RofA contains two putative DNA binding domains: the helix-turn helix mga domain (HTH_mga) (residues 7-65) and a mga DNA-binding domain (Mga) (76-156 residues), followed by a phosphotransferase system (PTS) regulation domain (PRD) of Mga located between residues 171-384 and a putative C-terminal PTS EIIB-like domain only identified by the structural analysis. The two adjacent RofA amino acid changes in M1_UK_ and related sublineages are in the PRD_mga domain (M318I and F319V) while the other amino acid change (D491N) is in the final section of a putative EIIB-like domain. *In silico* analysis suggests that, unlike other mga-like proteins, RofA may have just one longer fused functional PRD domain instead of two. PRD_*mga* domains are crucial for *mga* activation, through histidine phosphorylation events, in response to environmental sugar status, while EIIB-like domains may influence dimerization [11–14]. To determine if the 3 RofA SNPs between M1_global_ and M1_UK_ may have an impact on RofA histidine phosphorylation, 50 RofA model structures of each form were generated with the comparative modelling software MODELLER. Analysis highlighted a possible effect of the M318I substitution upon His278: The 50 superimposed M1_global_ models revealed that, in at least 3 different conformations, Met318 and His278 residues are close enough to produce steric hindrance (Figure 4), with a likely mode of interaction (His N to Met S) that is well described [15]. By contrast, no such interaction can occur in M1_UK_ as Met318 is replaced by an isoleucine residue, with a side chain comprising only non-polar carbon and hydrogen atoms (Figure 4). Consequently, the distribution of His278-M318I distances are larger in the M1_UK_ than in M1_global_ models (Figure S8), an effect that is further re-inforced by the shorter length of the Ile chain in M1_UK_.

**Figure 3.**
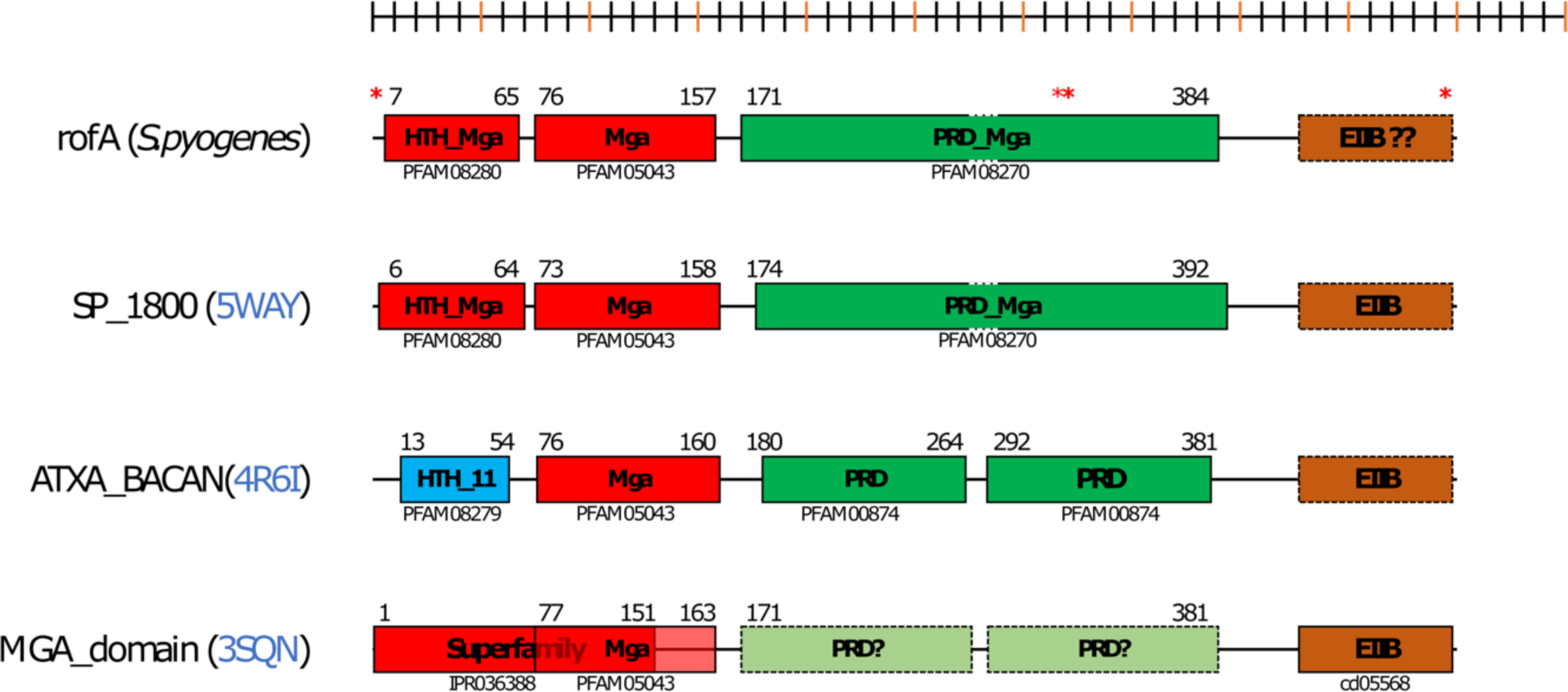
Schematic representation of RofA and *mga* homologous domain structures. Defined lines indicate domains that have been identified by their interpro signature. Blurred boundaries indicate domains predicted by structural similarity. Helix-turn-helix (HTH) and DNA-binding domains are represented in blue and red, phosphotransferase system regulatory domains (PRD) in green and putative EIIB domains in orange. From the top, diagram depicts RofA; MgaSpn, a Mga regulator from *Streptococcus pneumoniae* (5WAY); AtxA, a virulence regulator from *Bacillus anthracis* (4R6I); and putative Mga family transcriptional regulator from *Enterococcus faecalis* (3SQN).

**Figure 4.**
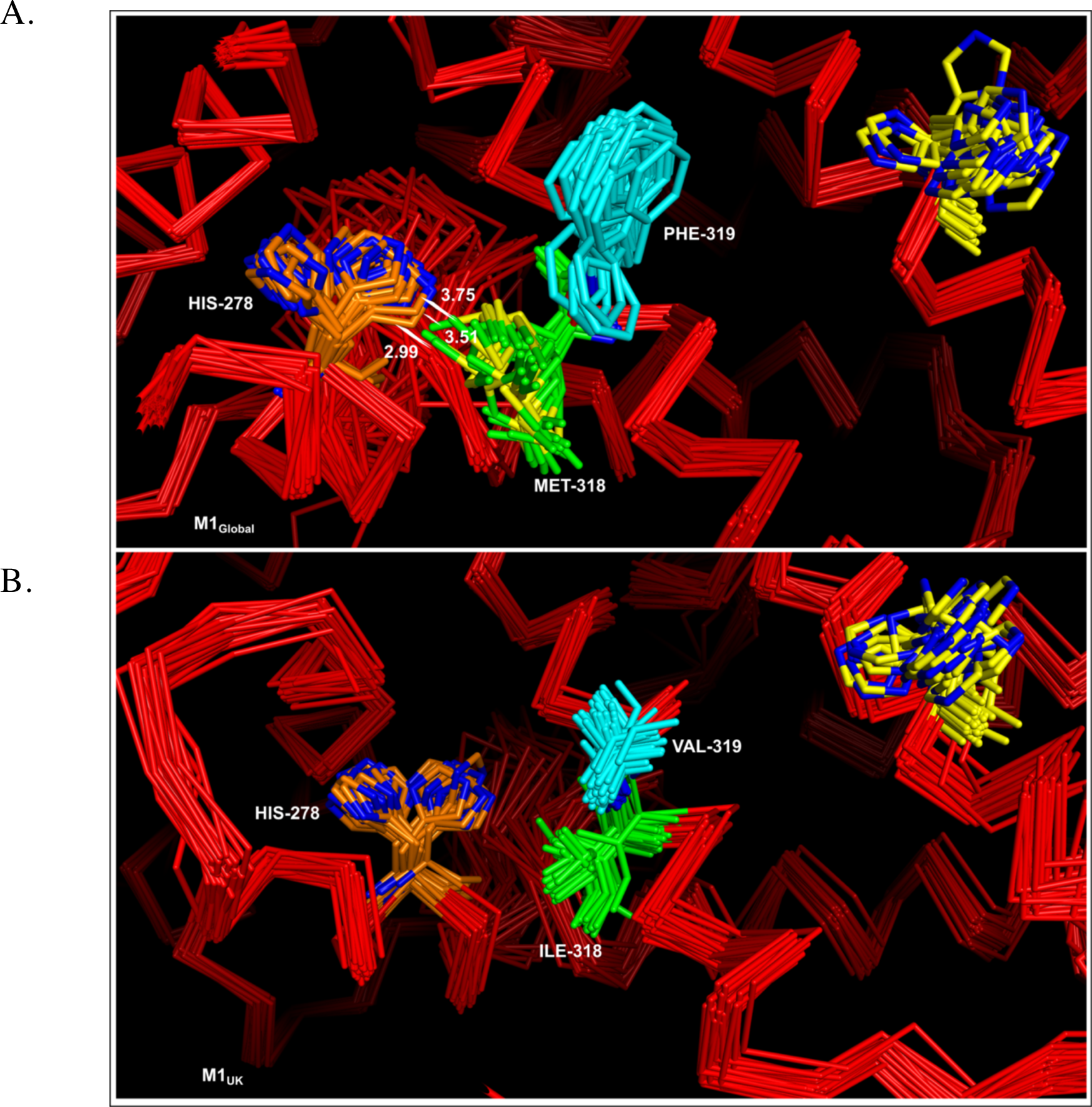
Conformations of His-278 and Met/Ile-318 in the 50 homology models of RofA in A) M1_global_ lineage strains; B) M1_UK_ lineage strains. His-278 is represented in orange, and Met/Ile318 residues in green. Interatomic distances between His278 and Met318 are represented by a line when they are below van der Waals contact distance for the corresponding atoms.

## Discussion

In this study, we analyzed the RofA regulon in 6 different *emm*1 strains, that were representative of the two contemporary major *emm*1 lineages (M1_global_ and M1_UK_). The transcriptome of RofA deletion mutants has not to our knowledge been fully reported; our results suggest that the transcriptomic changes may vary according to strain background, even within the same *emm* genotype. Although the pilus locus was a major target of RofA regulation in both lineages, RofA mutation led to a number of discrete transcriptomic changes that were unique to M1_UK_ strains and were not seen in M1_global_. Although introduction of the 3 SNPs into *rofA* made little or no impact in an M1_global_ strain, the reversal of 3 SNPs in an M1_UK_ strain led to an unexpected number of transcriptomic changes that, in part, recapitulated transcriptomic changes seen when deleting RofA. Computational analysis identified a key role for target residues in RofA that may interact with histidine residues, predicting that phosphorylation of RofA and function of the PRD domain may differ in M1_UK_ strains.

RofA was first identified as a positive regulator of the *prtF* genes in *emm*6 *S. pyogenes* [6]. Previous investigators have identified apparent serotype-specific differences in RofA target genes when evaluating *emm*2, *emm*6, and *emm*49 isogenic mutants; our data suggest that the range of target genes that are subject to RofA regulation may well vary from strain to strain despite sequence identity between promoters [2, 6, 16]. Based on our findings, the regulon of RofA cannot easily be explained by consensus DNA binding sequence motifs or involvement of other regulators. Indeed, the predicted motifs were quite different from previously published RofA motifs [17]. The sequence logos contained poly-AT tracts, which are known to be relatively flexible [18]. This suggests RofA may recognise DNA conformation or curvature, rather than a specific sequence, as proposed for the structurally-similar AtxA protein [19]. This would explain how this single protein can have such a variable regulon across the species, and even between closely-related isolates of the same *emm* type. Hence subtle changes to the protein’s structure, such as those caused by the three SNPs investigated in this study, may have an effect through differentially influencing RofA’s association with multiple loci, rather than simply changing its affinity for a clearly-defined binding motif.

We have undertaken systematic analysis of the RofA regulon in the widely successful *emm*1 pandemic lineage (M1_global_). Although our findings confirm the role of RofA as a strong positive regulator of the pilus locus in *emm*1 strains as previously reported, we also identified a clear role for RofA in regulation of Spy0212/0213, Spy1081(PTS system), Spy1281(two component regulator), phage protein (Spy1453) and *scpA*. RofA was reported to suppress transcription of *speA*, *sagA*, and *mga* [2] but we could not confirm this in *emm*1 strains. In the current study, RNAseq identified *speA* as a target for repression in only one *rofA* mutant. Although single M1_global_ and M1_UK_ strains showed upregulation of *sagA* upon *rofA* deletion, other strains from each lineage showed downregulation of *sagA*, underscoring the importance of considering more than one strain when examining regulatory gene control. When we compared the RofA regulon between M1_UK_ and M1_global_ we identified additional unique genes that were regulated in only M1_UK_ including *epf*, *glnQ.2*, the *bglA.2* operon, Spy1732 and Spy1736.

RNAseq data showed that 2 of the 3 M1_UK_ rofA mutant strains exhibited downregulation of the *ntpAB* operon that encodes V-type ATPases. Interestingly our recently reported proteomic data showed that strains from M1_UK_ and the M1_23SNP_ sublineage have reduced (rather than increased) levels of NtpA and B in the cytosol compared with strains from M1_global_ and the M1_13snp_ sublineage [3]. The role of the V-type ATPase in the physiology and pathogenesis of *S. pyogenes* is not known though the operon may be regulated by small RNAs [20]. V-type ATPases, located in the plasma membrane, couple the transfer of protons or sodium cations across the membrane with ATP hydrolysis or synthesis, and are responsible for cytoplasmic proton extrusion, regulating internal pH. [21] *S. pyogenes* performs only lactic acid fermentation for production of energy and can survive despite lowering the pH to ∼5.4 in growth medium [20]. Thus, the V-type ATPase in *S. pyogenes* might be involved in pumping hydrogen ions from the bacterial cytosol to overcome acid stress to survive the acidic conditions inside the host lysosome.

RofA also appears to play a role in positive regulation of the citrate lyase operon which is also involved in pH tolerance, in some, but not all, *emm*1 strains. One M1_global_ and two M1_UK_ rofA mutant strains showed reduced transcription in the *citABCDE* operon. Citrate lyase is a key enzyme that allows the microorganism to enter the citric acid cycle in the reductive mode and the citrate lyase operon may help *S. pyogenes* adapt to metabolic stress, such as low pH, lactate accumulation and nutrient-deprived, hypoxic conditions [22–24]. This metabolic “switch” facilitates survival during environmental transitions encountered in the infective process [24], while absence of the citrate lyase operon may render *emm* 12 *S. pyogenes* strains less fit under nutrient-deprived conditions [22].

The reversal of the 3SNPs in *rofA* in an M1_UK_ strain resulted in a transcriptomic effect that in part emulated *rofA* deletion in the same strain. This suggests that the function of *rofA* in M1_UK_ strains is somehow reliant on these 3SNPs, while this is not the case in M1_global_ strains, potentially because of the subsequent adaptive mutations that have occurred in M1_UK_. We noted upregulation of phage genes in the strain with reversal of the *rofA* 3SNPs that was unexpected, and may reflect a stress response. Davies et al recently reported the effects of reversal of the 3SNPs in an M1_UK_ strain, and found that no gene was differentially expressed by the repair of the *rofA* 3SNPs [4]. In contrast, we found 91 genes to be differentially regulated in the *rofA* repaired strain (10 genes downregulated and 81 genes up regulated). While this could be a strain-specific effect, or related to the different growth media used, or related to different expression thresholds used in each study, the data cannot be compared directly as the two studies used different reference strains.

Currently, the mechanism by which RofA is controlled is unknown. Experimental data relating to Mga (the closest related protein studied) suggests that the PRD domain is activated by histidine phosphorylation events that lead to a defect in protein oligomerization, altered gene expression and, in some cases, virulence attenuation [11–14]. The number and positions of phosphorylated histidines within PRD_*mga* domain vary among the regulators, and phosphorylation can positively or negatively affect protein activity [11]. Our protein model predictions show that the amino acid substitutions between M1_global_ and M1_UK_ may interfere with RofA histidine phosphorylation. In fact, M318I replacement could disrupt a stabilizing interaction between this residue and His278. This native interaction could bias the His conformation towards the proximal rotamer (when the His residue is nearest to M318), limiting access to its kinase enzyme and resulting in different phosphorylation activation events between lineages. Whether the PRD domain of RofA is indeed phosphorylated at His278 is currently unknown, as is the experimental impact of the 3SNPs on phosphorylation of RofA.

Our study has highlighted the importance of examining multiple strains when considering *S. pyogenes* regulators, however a limitation is the number of strains that can be practically examined in different growth conditions. This extends to the testing of strains *in vivo*, where comparison of multiple isogenic strains or different lineages in the same model can be challenging and may require a prohibitive number of mice per group to demonstrate a clear difference. The acquisition of the 3SNPs in *rofA* in the M1T1 *S. pyogenes* lineage is unique to the *emm*1 intermediate sublineages and M1_UK_; we have not detected these 3SNPs in any other *emm* genotype. As such, the SNPs act as a useful marker of the new lineages. The 3SNPs in *rofA* were identified as early as 2005 [3] and have persisted throughout the evolution of M1_UK_, present in both M1_13SNP_ and M1_23SNP_ strains, underlining a likely key role in the success of the novel lineage.

## Materials and methods

### Ethics statement

All animal use was licensed by a U.K. Home Office Project Licence using protocols approved by the local ethical review process. Whole heparinised human blood from consenting normal donors from a subcollection of the Imperial Tissue Bank (ICHTB, approved by Wales REC reference 17/WA/0161)

### Bacterial strains and growth conditions

*S. pyogenes* strains were all non-invasive clinical isolates that were collected for the purpose of sequential genotyping and genome sequencing. Strains used in this study are listed in Supplementary Table S1. Routinely, *S. pyogenes* were cultured on Columbia blood agar (CBA) (OXOID, Basingstoke, UK), Todd-Hewitt (TH) agar or in TH broth (OXOID) at 37°C with 5% CO_2_ for 16 hours. Chemically defined medium (CDM) supplemented with different sugars was also used for growth of *S. pyogenes*. Where appropriate, spectinomycin (50 μg/ml) or kanamycin (400 μg/ml) were added to the culture medium for *S. pyogenes*. *Escherichia coli* strains Top10 (Invitrogen) and DH5α were used for cloning and were grown in Luria broth (LB) or on Luria broth agar with kanamycin (50 μg/ml) or spectinomycin (50 μg/ml).

Isolates for RNASeq analysis were selected from genome-sequenced non invasive *S. pyogenes* isolates (Supplementary Table S1). The inclusion criteria were that isolates were either in the M1_global_ or in the M1_UK_ lineages not intermediates; possessed a phage containing the *speA* gene; were wildtype for *covRS;* with overall phage and superantigen content matching the reference *emm*1 sequence MGAS5005.

### Whole blood bacterial survival assay

Whole heparinised human blood was used in Lancefield assays as previously described [25]. Briefly approximately 50 CFU of *S. pyogenes* were inoculated into 300 μL whole blood and incubated at 37°C, rolling for 3 hours; each strain was tested in 3 normal donor whole blood samples (with technical triplicates for each donor). Input and output colony forming units were measured by plating and the multiplication factor calculated.

### Construction of a *rofA* disruption mutant

A 491 bp fragment downstream of the *rofA* gene was amplified (forward primer: *5′-GGAATTCCTCTTACATAAGATTCATATC -3′*, reverse primer: *5′-GGGGTACCCTCTTCC-TACACTTAGAAAGC -3′*) incorporating EcoRI and KpnI restriction sites into the 5′ and 3′ ends respectively, and cloned into the suicide vector pUCMUT to produce vector pUCMUT_rofAD_. A 523 bp fragment upstream of the *rofA* gene was amplified (forward primer: *ACGCGTCGAC-CGCCATGTCACCACATTGCG -3′*, reverse primer: *5′-AACTGCAGGGGTTACCTGTGCCA-TAATC -3′*) incorporating PstI and SalI restriction sites into the 5′ and 3′ ends respectively and cloned into PstI/SalI digested pUCMUT_rofAU_. The construct was introduced into M1_global_ strains (H1488, H1489 and H1499) and MI_UK_ strains (H1491, H1496, and H1490) by electroporation and crossed into the chromosome by homologous recombination. Transformants were selected using kanamycin (400μg/ml). Successful disruption of the *rofA* gene and insertion of the kanamycin resistance cassette was confirmed by PCR, DNA sequencing, and whole genome sequencing.

### Complementation of *rofA* strains

The *rofA* coding sequence, including native promoter, was amplified from both M1_global_ and M1_UK_ *S. pyogenes* DNA (forward primer: *5′-GACGCATGCCTCCTCTCAATGTGACATC-3′*, reverse primer: *5′-ACGGATCCGTGGTGACATGGCGCTTATGTT-3′*) incorporating BamHI and SphI restriction sites to each end of the PCR products, and cloned into BamHI and SphI digested shuttle vector pDL278 resulting in plasmid pDL_rofAM1_ and pDL_rofAM1UK_, which were confirmed by Sanger sequencing. The resulting plasmids were introduced into M1_globalrofAKO_ (H1561) and M1_UKrofAKO_ (H1582) by electroporation. The successful introduction of plasmid was confirmed by PCR specific for pDL278 backbone (forward primer: *5′-CATTCAGGCTGCGCAACTG-3′*, reverse primer: *5′-TCGAATTCACTGGCCGTCG-3′*) in each of the resulting isogenic strains H1591 (M1_globalrofAKO_ complementation) and H1592 (M1_UKrofAKO_ complementation).

### Construction of *rofA* 3SNP mutant strains

To introduce the 3SNPs of *rofA* typical of M1_UK_ into M1_global_ (H1488), a 597 bp fragment downstream of the *rofA* gene was amplified (forward primer: *5′-* AACTGCAGAG-CACATTAAGTCCGATTGCAG*-3′*, reverse primer: *5′-* ACGCGTCGACCCACAC-CTTAACTTAATCCCGA*-3′*) incorporating *EcoRI* and *KpnI* restriction sites into the 5′ and 3′ ends respectively, and cloned into the suicide vector pUCMUT to produce vector pUCMUT_ro-fAD_. A 1548 bp fragment upstream of the *rofA* gene was amplified (forward primer: *5′-* GGGG-TACCCTAAAGTCGCGCAATGTGGTG*-3′*, reverse primer: *5′-* AGCGAATTCGTGTAG-GAAGAGAGGTCCCT *-3′*) incorporating *PstI* and *SalI* restriction sites into the 5′ and 3′ ends respectively and cloned into *PstI*/*SalI* digested pUCMUT_rofAU_. The construct was introduced into M1_global_ (H1488) by electroporation and crossed into the chromosome by homologous re-combination. Transformants were selected using kanamycin (400μg/ml). Successful allelic replacement and insertion of the kanamycin resistance cassette was confirmed by PCR, DNA sequencing and whole genome sequencing.

To fix the *rofA* 3SNPs in M1_UK_ (H1496), a 597 bp fragment downstream of the *rofA* gene was amplified (forward primer: *5′-* AACTGCAGAGCACATTAAGTCCGATTGCAG*-3′*, reverse primer: *5′-* ACGCGTCGACCCACACCTTAACTTAATCCCGA *-3′*) incorporating *EcoRI* and *KpnI* restriction sites into the 5′ and 3′ ends respectively and cloned into the suicide vector pUCMUT to produce vector pUCMUT_rofAD_. A 1548bp fragment of the *rofA* gene was amplified (forward primer: GGGGTACCCTAAAGTCGCGCAATGTGGTG *-3′*, reverse primer: *5′-* AGCGAATTCGTGTAGGAAGAGAGGTCCCT *-3′*) incorporating *PstI* and *SalI* restriction sites into the 5′ and 3′ ends respectively and cloned into *PstI*/*SalI* digested pUCMUT_rofAU_. The construct was introduced into M1_UK_ (H1496) by electroporation and crossed into the chromosome by homologous recombination. Transformants were selected using Kanamycin (400μg/ml). Successful allelic replacement and insertion of the kanamycin resistance cassette was confirmed by PCR and DNA and whole genome sequencing.

### Quantitative real-time PCR

The extraction of RNA was done as described previously [26]. *S. pyogenes* were grown in THY broth until late logarithmic growth phase. The bacterial cultures were treated with aqua-phenol and phenol: chloroform: isoamyl alcohol (Sigma, UK), and then precipitated with 2-propanol, followed by DNase treatment with TurboDNAfree (Ambion, Cambridgeshire UK). RNA was reverse transcribed into complementary DNA (cDNA) using Transcriptor reverse transcriptase kit (Roche Diagnostics, UK). qRT-PCR was carried out for Spy0107, Spy0109, and *speA* using a Real time PCR System (Thermo Fisher Scientific) [3] and expression data normalized to that of housekeeping gene *proS* using a standard curve method as described previously [27].

### Transcriptome (RNA-Seq) analysis

*S. pyogenes* were grown in THY broth until late logarithmic growth phase. 3 M1_global_ vs 3 M1_global RofAKO_, 3 M1_UK_ vs 3 M1_UKRofAKO_ were grown in triplicate (Table S1), RNA was prepared as previous described [26], and the quality and quantity of total RNA was evaluated using an Agilent 2100 Bioanalyzer with the RNA 6000 Nano Total RNA Kit. Ribosomal RNA was depleted using NEBNext^®^ rRNA Depletion Kit (Bacteria), and RNAseq NGS libraries made using the NEBNext® Ultra™ II Directional RNA Library Prep Kit for Illumina® according to manufacturer’s instructions, by the MRC Genomics Core Lab. A minimum of 10 million paired end 100bp reads were generated for each sample on HiSeq 2500 and NextSeq 2000 Illumina Sequencers. Illumina sequencing data have been submitted to the European Nucleotide Archive (ENA, www.ebi.ac.uk/ena) project PRJEB62819 with the accession numbers given in Table S1. RNA-seq data were analyzed according to the followed pipeline. Read quality was accessed using FastQC (https://www.bioinformatics.babraham.ac.uk/projects/fastqc/), filtered and trimmed using trimmomatic [27], and mapped against the MGAS5005 (CP000017) reference genome using bowtie2 [28], with the highest sensitivity options. The resulting alignments were converted to sorted BAM files using vcftools [29]. Initial visualizations of the sequencing mapping were performed using the Integrative Genomics Viewer (IGV) [30], including the confirmation of *rofA* disruptions, 3 SNPs insertions, and *emm*1 lineages. The mapped RNA-seq reads were then transformed into a fragment count per gene per sample using HT-seq [31] packages and featureCounts [32] and the main results compared. Exploratory data analysis (Principal component analysis and Heatmap of sample-to-sample distances) of the RNAseq data was implemented and plotted using DESeq2 package [33]. Differential expression analysis in each dataset was performed using three different R packages (DESeq2 [33], EdgeR [34] and limma (https://bioconductor.riken.jp/packages/3.0/bioc/html/limma.html)) with a log2fold change of 0.5 and p-value < 0.01 and p-adj < 0.05. Only genes DE in two of the three software’s used were considered as DE genes and used in the following analysis. Prophage regions were predicted using phaster [35], and curated by visual assessment and blast alignment. Operon prediction was performed using SP119 annotation [36] and the motifs analysis was performed using XTREME (https://meme-suite.org/meme/doc/xstreme.html). Correlation coefficients for RNA-seq were determined by plotting the log2 value of the array on the x axis to the log2 value of the quantitative real-time PCR on the y axis. Linear regression was used to determine the line of best fit, and the resulting R2 value was calculated, which represented the fitness of the data.

### Murine intra-nasal infection

FVB/n female mice 6–8 weeks old (Charles River, Margate, UK) were briefly anaesthetized with isofluorane and challenged intra-nasally with 5×10^6^ CFU *S. pyogenes*, administered as 5µl per nostril. Nasal shedding was longitudinally and non-invasively monitored daily for 7 days following intranasal challenge using a nose-pressing technique [37]. Briefly, murine noses were pressed gently onto a CBA plate (Oxoid) 10 times, every 24h. Resulting exhaled moisture was spread over the plate and colonies counted following incubation at 37°C, 5% CO_2_ for 24 hours. On day 7 mice were euthanized, and nasal tissues, spleen and liver dissected and plated onto CBA to quantify nasal and systemic *S. pyogenes* burden. Isogenic strains were compared over 7 days. To detect airborne shedding of bacteria within cages of infected mice, CBA settle plates (4 per cage) were placed face up on the upper rack of individually HEPA filtered cages for 4 hours on days, 0, 1, 2 and 3 post-infection as previously reported [37]. Airborne dispersal of *S. pyogenes* was quantified, by counting beta haemolytic colonies following overnight incubation of plates at 37°C, 5% CO_2_.

### Comparative model building and histidine location predictions

No homologues of Spy0106 *rofA* were found in Protein Data bank (PDB) sequences of known structure. Therefore, more sensitive methods for the detection of weak homologues were required. Two different approaches were used: an Hiden Markov Model (HMM) based search using HH pred [38] and the threading server LOMETS (Zhang Lab) [39], both consistently leading to the identification of three PDB sequences (PDB codes 4R6I, 5WAY and 3SQN). The signature motifs identified in the three proteins were confirmed with a HMMER search [40]. Structural alignment and search of the three identified structures using the FATCAT flexible alignment server [41] confirmed their homology at the structural level, which suggests that any of them could be used as a template for model generation. 4R6I (AtxA protein, avirulence regulator from *Bacillus anthracis*) was the chosen template, as it had the longest alignment coverage of the target and the highest significance for the detected motifs. Using this template, models for M1_rofAKO_ and M1_UKrofAKO_ were produced using the MODELLER software [42]. The MODELLER template-target (guide) alignment was generated with the HHPRED server, and 50 models were produced for each of the *rofA* variants, using the recommended MODELLER parameters for a thorough structure search and minimization (MODELLER Manual, v 10.1). The quality of the generated models was assessed with the Ga341 score [43], the QMEAN score [44] and the MOLPROBITY server [45]. Each set of 50 models was loaded and visualized in the molecular viewer software PyMOL (Schrodinger LLC) to access structural variance and the potential impact of amino acid substitutions between variants.

### Statistical analysis

All statistical analyses were performed with GraphPad Prism 5.0. Comparison of two datasets was carried out using unpaired students t-test and three or more data sets were analysed by Kruskal-Wallis followed by Dunn’s multiple comparison test or ANOVA and Bonferroni post-test depending on sample size. Survival data were analysed by Mantel-Cox (Log rank) test. A p-value of ≤0.05 was considered significant.

## Acknowledgements

The authors acknowledge the support of the NIHR Imperial Biomedical Research Centre and Colebrook Laboratory in collection and curation of clinical *S. pyogenes* throat isolates and the Imperial College Healthcare Trust Tissue bank for access to normal donor blood samples. They also thank the MRC LMS sequencing facility for undertaking RNA sequencing during the early stages of the COVID-19 pandemic and acknowledge support from UKHSA colleagues Drs Theresa Lamagni and Vicki Chalker in securing funding for work on M1_UK_. HKL is an MRC CMBI Clinical Training Fellow and NC is a Sir Henry Dale Fellow, jointly funded by Wellcome and the Royal Society (grant no. 104169/Z/14/A).

## References

1. Lynskey NN, Jauneikaite E, Li HK, Zhi X, Turner CE, Mosavie M, et al. Emergence of dominant toxigenic M1T1 *Streptococcus pyogenes* clone during increased scarlet fever activity in England: a population-based molecular epidemiological study. The Lancet Infectious Diseases. 2019;19(11):1209–18.

2. Beckert S, Kreikemeyer B, Podbielski A. Group A streptococcal rofA gene is involved in the control of several virulence genes and eukaryotic cell attachment and internalization. Infect Immun. 2001;69(1):534–7. Epub 2000/12/19. doi: 10.1128/IAI.69.1.534-537.2001.

3. Li HK, Zhi X, Vieira A, Whitwell HJ, Schricker A, Jauneikaite E, et al. Characterization of emergent toxigenic M1_UK_ *Streptococcus pyogenes* and associated sublineages. Microbial Genomics. 2023;9(4).

4. Davies MR, Keller N, Brouwer S, Jespersen MG, Cork AJ, Hayes AJ, et al. Detection of *Streptococcus pyogenes* M1(UK) in Australia and characterization of the mutation driving enhanced expression of superantigen SpeA. Nat Commun. 2023;14(1):1051. Epub 20230224. doi: 10.1038/s41467-023-36717-4.

5. Abbot EL, Smith WD, Siou GP, Chiriboga C, Smith RJ, Wilson JA, et al. Pili mediate specific adhesion of *Streptococcus pyogenes* to human tonsil and skin. Cellular Microbiology. 2007;9(7):1822–33.

6. Fogg GC, Gibson CM, Caparon MG. The identification of rofA, a positive-acting regulatory component of prtF expression: use of an mγδ-based shuttle mutagenesis strategy in *Streptococcus pyogenes*. Molecular Microbiology. 1994;11(4):671–84.

7. Granok AB, Parsonage D, Ross RP, Caparon MG. The RofA binding site in *Streptococcus pyogenes* is utilized in multiple transcriptional pathways. Journal of Bacteriology. 2000;182(6):1529–40.

8. Kreikemeyer B, McIver KS, Podbielski A. Virulence factor regulation and regulatory networks in *Streptococcus pyogenes* and their impact on pathogen–host interactions. Trends in Microbiology. 2003;11(5):224–32. doi: 10.1016/s0966-842x(03)00098-2.

9. Kratovac Z, Manoharan A, Luo F, Lizano S, Bessen DE. Population genetics and linkage analysis of loci within the FCT region of *Streptococcus pyogenes*. J Bacteriol. 2007 Feb;189(4):1299–310. doi: 10.1128/JB.01301-06.

10. Calfee G, Danger JL, Jain I, Miller EW, Sarkar P, Tjaden B, et al. Identification and characterization of serotype-specific variation in group A Streptococcus pilus expression. Infection and Immunity. 2018;86(2):e00792–17.

11. Zeng S, Liu Y, Wu M, Liu X, Shen X, Liu C, et al. Identification and validation of reference genes for quantitative real-time PCR normalization and its applications in lycium. PloS One. 2014;9(5):e97039.

12. Hondorp ER, Hou SC, Hempstead AD, Hause LL, Beckett DM, McIver KS. Characterization of the Group A Streptococcus Mga virulence regulator reveals a role for the C-terminal region in oligomerization and transcriptional activation. Molecular Microbiology. 2012;83(5):953–67.

13. Hondorp ER, McIver KS. The Mga virulence regulon: infection where the grass is greener. Molecular Microbiology. 2007;66(5):1056–65.

14. Tsvetanova B, Wilson AC, Bongiorni C, Chiang C, Hoch JA, Perego M. Opposing effects of histidine phosphorylation regulate the AtxA virulence transcription factor in *Bacillus anthracis*. Molecular Microbiology. 2007;63(3):644–55.

15. Pal D, Chakrabarti P. Non-hydrogen bond interactions involving the methionine sulfur atom. Journal of Biomolecular Structure and Dynamics. 2001;19(1):115–28.

16. Podbielski A, Woischnik M, Leonard BA, Schmidt KH. Characterization of nra, a global negative regulator gene in group A streptococci. Molecular Microbiology. 1999;31(4):1051–64.

17. Fogg GC, Caparon MG. Constitutive expression of fibronectin binding in *Streptococcus pyogenes* as a result of anaerobic activation of rofA. Journal of Bacteriology. 1997;179(19):6172–80.

18. Gordon BR, Li Y, Cote A, Weirauch MT, Ding P, Hughes TR, Navarre WW, Xia B, Liu J. Structural basis for recognition of AT-rich DNA by unrelated xenogeneic silencing proteins. Proc Natl Acad Sci U S A. 2011 Jun 28;108(26):10690–5. doi: 10.1073/pnas.1102544108.

19. Hadjifrangiskou M, Koehler TM. Intrinsic curvature associated with the coordinately regulated anthrax toxin gene promoters. Microbiology (Reading). 2008 Aug;154(Pt 8):2501–2512. doi: 10.1099/mic.0.2007/016162-0.

20. Tesorero RA, Yu N, Wright JO, Svencionis JP, Cheng Q, Kim J-H, et al. Novel regulatory small RNAs in *Streptococcus pyogenes*. PloS One. 2013;8(6):e64021.

21. Magalhães PP, Paulino TP, Thedei Jr G, Ciancaglini P. Kinetic characterization of P-type membrane ATPase from *Streptococcus mutans*. Comparative Biochemistry and Physiology Part B: Biochemistry and Molecular Biology. 2005;140(4):589–97.

22. Vlaminckx BJ, Schuren FH, Montijn RC, Caspers MP, Fluit AC, Wannet WJ, et al. Determination of the relationship between group A streptococcal genome content, M type, and toxic shock syndrome by a mixed genome microarray. Infection and Immunity. 2007;75(5):2603–11.

23. Martín MG, Sender PD, Peirú S, De Mendoza D, Magni C. Acid-inducible transcription of the operon encoding the citrate lyase complex of *Lactococcus lactis biovar diacetylactis* CRL264. Journal of Bacteriology. 2004;186(17):5649–60.

24. Srinivasan V, Morowitz HJ. Ancient genes in contemporary persistent microbial pathogens. The Biological Bulletin. 2006;210(1):1–9.

25. Lynskey NN, Goulding D, Gierula M, Turner CE, Dougan G, Edwards RJ, et al. RocA truncation underpins hyper-encapsulation, carriage longevity and transmissibility of serotype M18 group A streptococci. PLoS Pathog. 2013;9(12):e1003842. Epub 2013/12/25. doi: 10.1371/journal.ppat.1003842.

26. Turner CE, Kurupati P, Jones MD, Edwards RJ, Sriskandan S. Emerging role of the interleukin-8 cleaving enzyme SpyCEP in clinical *Streptococcus pyogenes* infection. Journal of Infectious Diseases. 2009;200(4):555–63.

27. Bolger AM, Lohse M, Usadel B. Trimmomatic: a flexible trimmer for Illumina sequence data. Bioinformatics. 2014;30(15):2114–20.

28. Clausen PT, Aarestrup FM, Lund O. Rapid and precise alignment of raw reads against redundant databases with KMA. BMC Bioinformatics. 2018;19:1–8.

29. Danecek P, Auton A, Abecasis G, Albers CA, Banks E, DePristo MA, et al. The variant call format and VCFtools. Bioinformatics. 2011;27(15):2156–8.

30. Robinson JT, Thorvaldsdóttir H, Winckler W, Guttman M, Lander ES, Getz G, et al. Integrative genomics viewer. Nature Biotechnology. 2011;29(1):24–6.

31. Anders S, Pyl PT, Huber W. HTSeq—a Python framework to work with high-throughput sequencing data. Bioinformatics. 2015;31(2):166–9.

32. Liao Y, Smyth GK, Shi W. featureCounts: an efficient general purpose program for assigning sequence reads to genomic features. Bioinformatics. 2014;30(7):923–30.

33. Love M, Anders S, Huber W. Differential analysis of count data–the DESeq2 package. Genome Biol. 2014;15(550):10–1186.

34. Robinson MD, McCarthy DJ, Smyth GK. edgeR: a Bioconductor package for differential expression analysis of digital gene expression data. Bioinformatics. 2010;26(1):139–40.

35. Arndt D, Grant JR, Marcu A, Sajed T, Pon A, Liang Y, et al. PHASTER: a better, faster version of the PHAST phage search tool. Nucleic Acids Research. 2016;44(W1):W16–W21.

36. Rosinski-Chupin I, Sauvage E, Fouet A, Poyart C, Glaser P. Conserved and specific features of *Streptococcus pyogenes* and *Streptococcus agalactiae* transcriptional landscapes. BMC Genomics. 2019;20:1–15.

37. Alam FM, Turner CE, Smith K, Wiles S, Sriskandan S. Inactivation of the CovR/S virulence regulator impairs infection in an improved murine model of *Streptococcus pyogenes* naso-pharyngeal infection. PLoS One. 2013;8(4):e61655.

38. Hildebrand A, Remmert M, Biegert A, Söding J. Fast and accurate automatic structure prediction with HHpred. Proteins: Structure, Function, and Bioinformatics. 2009;77(S9):128–32.

39. Wu S, Zhang Y. LOMETS: a local meta-threading-server for protein structure prediction. Nucleic Acids Research. 2007;35(10):3375–82.

40. Potter SC, Luciani A, Eddy SR, Park Y, Lopez R, Finn RD. HMMER web server: 2018 update. Nucleic Acids Research. 2018;46(W1):W200–W4.

41. Li Z, Jaroszewski L, Iyer M, Sedova M, Godzik A. FATCAT 2.0: towards a better understanding of the structural diversity of proteins. Nucleic Acids Research. 2020;48(W1):W60–W4.

42. Martí-Renom MA, Stuart AC, Fiser A, Sánchez R, Melo F, Šali A. Comparative protein structure modeling of genes and genomes. Annual Review of Biophysics and Biomolecular Structure. 2000;29(1):291–325.

43. Melo F, Sali A. Fold assessment for comparative protein structure modeling. Protein Sci. 2007 Nov;16(11):2412–26. doi: 10.1110/ps.072895107

44. Benkert P, Biasini M, Schwede T. Toward the estimation of the absolute quality of individual protein structure models. Bioinformatics. 2011;27(3):343–50.

45. Williams CJ, Headd JJ, Moriarty NW, Prisant MG, Videau LL, Deis LN, et al. MolProbity: More and better reference data for improved all-atom structure validation. Protein Science. 2018;27(1):293–315.

