## Supplementary Figures S1-S8 for "Characterization of the RofA regulon in the pandemic M1_global_ and emergent M1_UK_ lineages of *Streptococcus pyogenes*"



| | |
|---|---|
| Figure S1. | Transcript copy number for pilus locus genes *spy0107* and *spy0109* |
| Figure S2 | Comparison of absolute transcript copy number for *SpeA* |
| Figure S3 | *S. pyogenes* growth in chemically defined medium. |
| Figure S4 | Multiplication factor of *S. pyogenes* in whole human blood. |
| Figure S5 | *S. pyogenes* M1$_{global}$ isogenic strains nasopharyngeal carriage and shedding |
| Figure S6. | Predicted RofA motifs in each dataset compared with Granok et al 2000 |
| Figure S7 | *S. pyogenes* M1$_{UK}$ isogenic strains nasopharyngeal carriage and shedding |
| Figure S8. | Distribution of minimum interatomic distances between His-278 and Met-318 in M1$_{global}$ and M1$_{UK}$ models |

**Figure S1.**

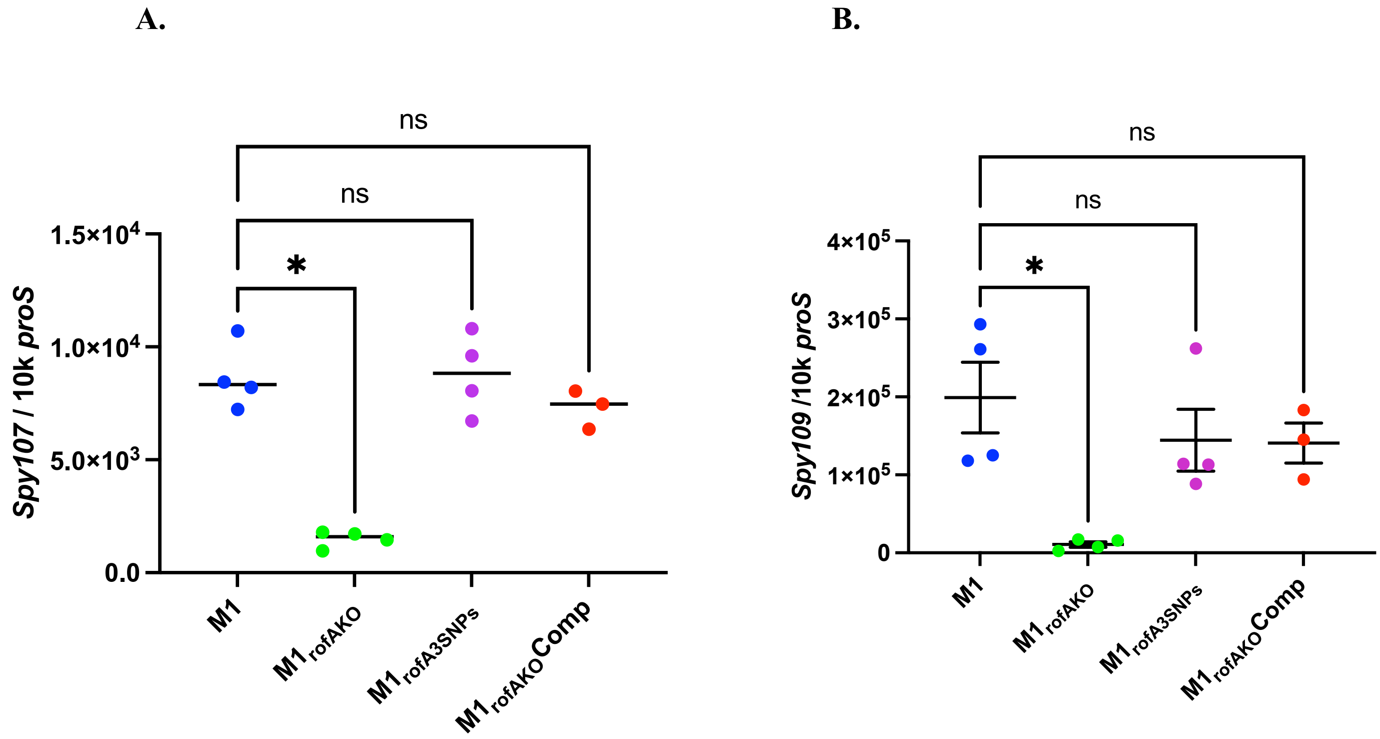


**Supplementary Figure S1. Comparison of absolute transcript copy number for pilus locus genes *spy0107* (A) and *spy0109* (B) standardized to housekeeping gene *proS*.** Comparison is between parent $M1_{global}$ strain H1488 ($M1_{global}$) with isogenic derivatives. Bars represent the mean and SEM of 4 biological replicates; each replicate shown as a coloured dot (average of technical triplicates). ( *=$p<0.05$).

**Figure S2**

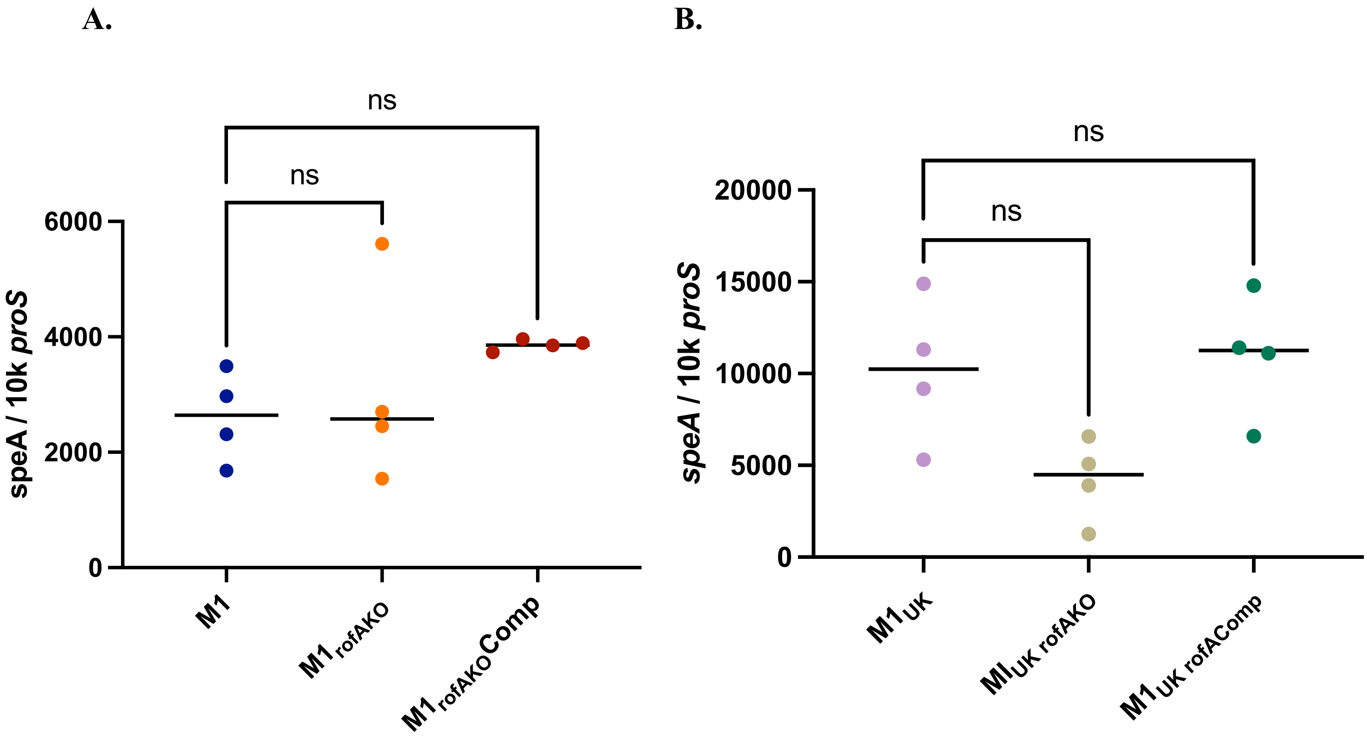


**Supplementary Figure S2. Comparison of absolute transcript copy number for *SpeA*** standardized to housekeeping gene *proS* in isogenic derivatives of $M1_{global}$ (A) and $M1_{UK}$ (B). Parent strains are H1488 ($M1_{global}$) and H1496 ($M1_{UK}$). Bars represent the mean and SEM of 4 biological replicates; each replicate shown as a coloured dot (average of technical triplicates). ns, not significant.

**Figure S3.**

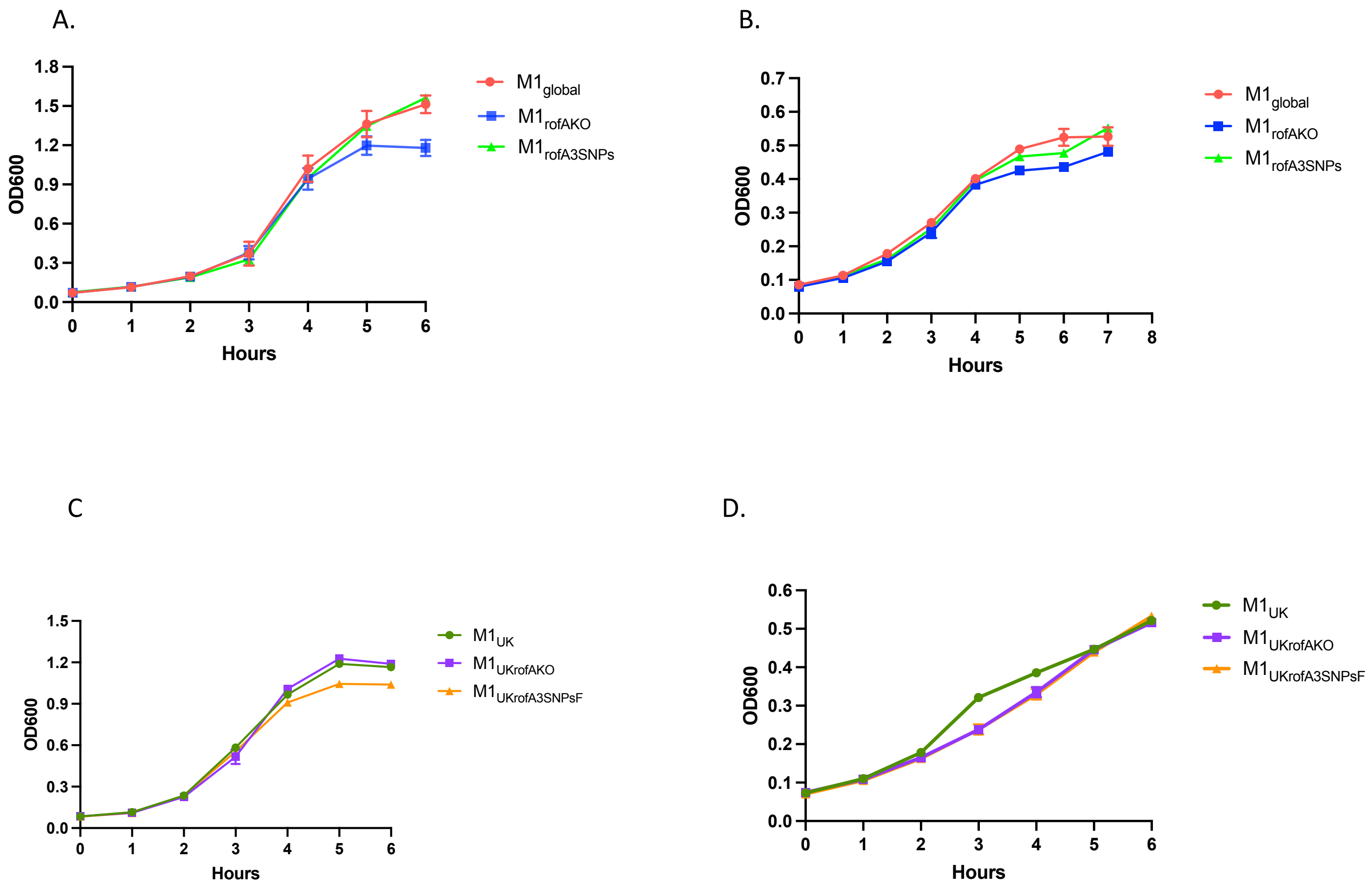


**Supplementary Figure S3. *S. pyogenes* growth in chemically defined medium.** Isogenic derivatives of M1$_{\mathrm{global}}$ strain H1488 (A and B) and M1$_{\mathrm{UK}}$ strain H1496 (C and D) are shown. M1$_{\mathrm{global}}$, M1$_{\mathrm{rofAKO}}$ and M1$_{\mathrm{rofA3SNPs}}$ strains supplemented with glucose (A) and mannose (B). M1$_{\mathrm{UK}}$, M1$_{\mathrm{UKrofAKO}}$ and M1$_{\mathrm{UKrofA3SNPsFixed\ (F)}}$ strains supplemented with glucose (C) and mannose (D). Data represent mean and SEM of three cultures at each time point.

**Figure S4**

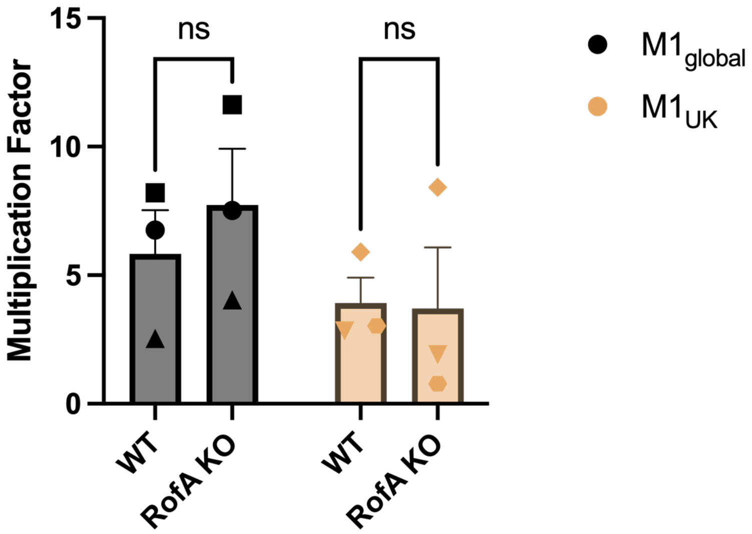


**Supplementary Figure S4. Multiplication factor of *S. pyogenes* in whole human blood.** Bars represent overall mean and SEM of three $M1_{global}$ or three $M1_{UK}$ strains, compared to the respective isogenic *rofA* deletion strains (n=3 pairs per lineage). Each point represents the mean multiplication factor (technical triplicates) of each individual strain tested in the blood of three healthy donors. Individual pairs of isogenic mutants are indicated by shape of dot. Note overall low multiplication factor.

**Figure S5**

**A.**

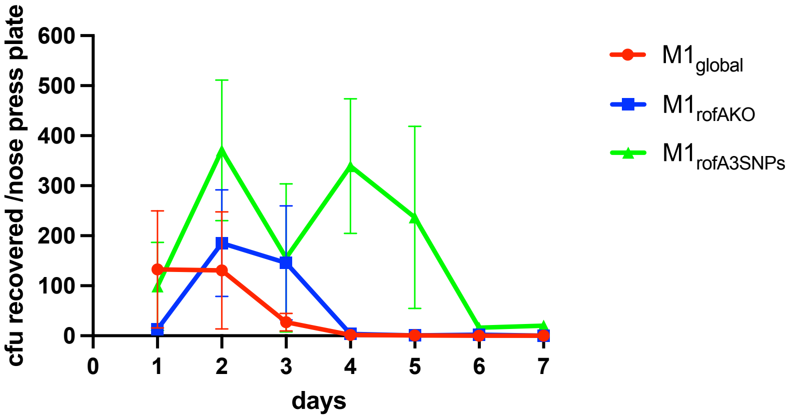


**B.**

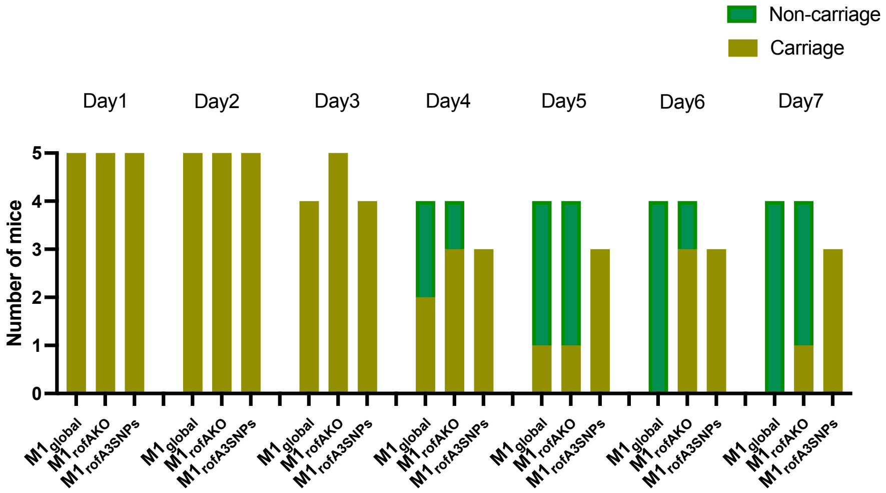


**C.**

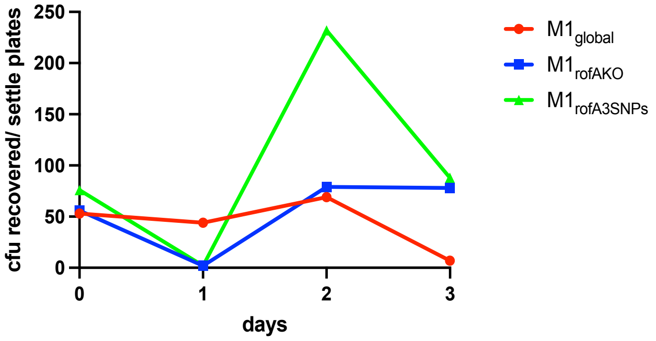


**Supplementary Figure S5.** ***S. pyogenes*** **$M1_{global}$ isogenic strains nasopharyngeal carriage and shedding.** Quantification of *S. pyogenes* nasal shedding as mean number of CFU recovered by daily nose pressing onto individual blood agar plates (A) and longevity of nasal shedding (B) comparing infection with isogenic $M1_{global}$, $M1_{rofAKo}$ and $M1_{rofA3SNPs}$ strains (n= 5 per group). Note reduction in group size day 3 onwards. Quantification of *S. pyogenes* shedding onto settle plates on days 0-3 of infection (n=4 plates/cage) as metric of shedding into air (C).

### Figure S6

A.

B.

C.

D.

E.

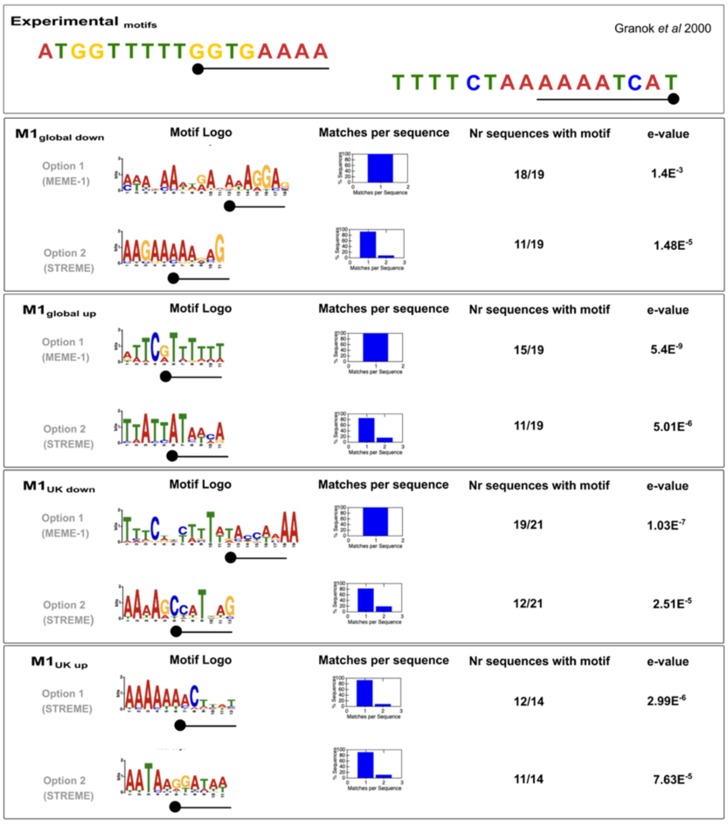


**Supplementary Figure S6.** Predicted RofA motifs in each dataset compared with motif experimentally identified by Granok et al 2000 (reference 7) (A). The motifs predicted by MEME-1 and/or STREME (B-E) are represented by position-specific probability matrices that specify the probability of each possible letter appearing at each possible position in an occurrence of the motif. The sequence LOGOS contain stacks of letters at each position in the motif. The height of the individual letters in a stack is the probability of the letter at that position multiplied by the total information content of the stack. In each dataset, the intergenic region of DE genes in two of the three strains sequenced was used. In the case that genes were in operons, according to Rosinski-Chupin 2019 (reference 36), the intergenic region of the operon was used. For $M1_{global}$ downregulated genes (B) were considered separately from upregulated genes (C). Similarly, for $M1_{UK}$, downregulated genes (D) were separated from upregulated genes (E).

**Figure S7**

A.

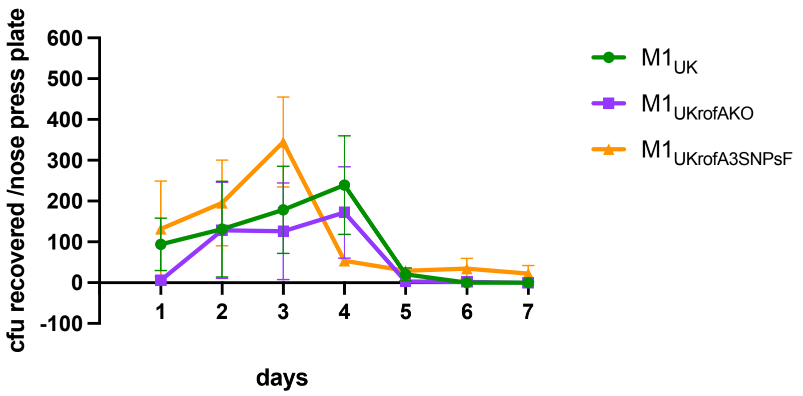


B.

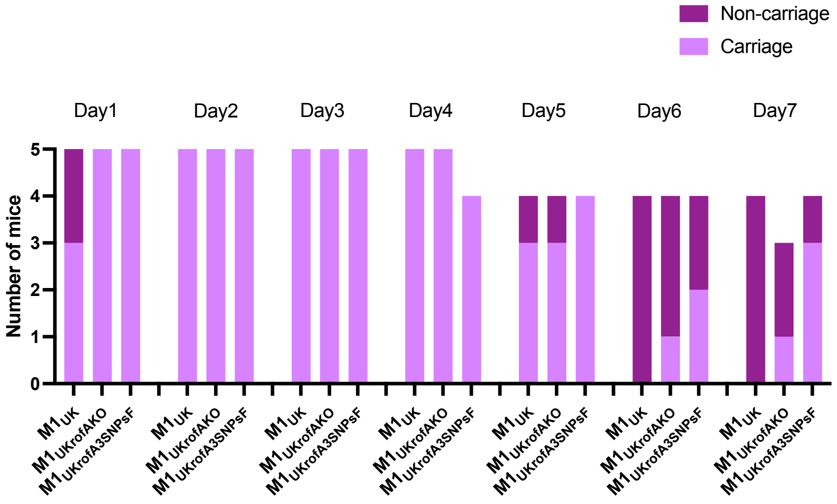


C.

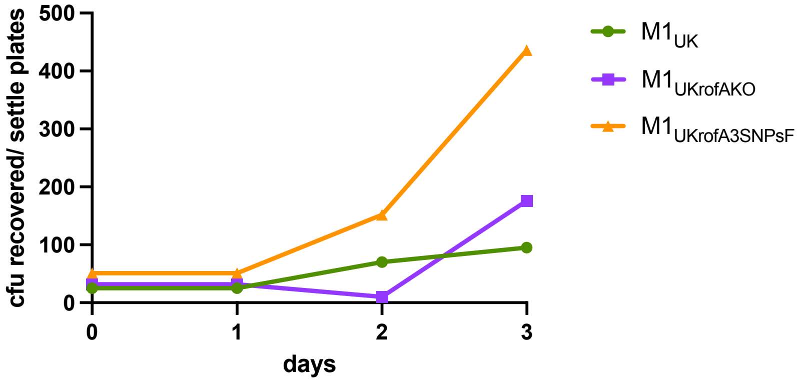


**Supplementary Figure S7.** ***S. pyogenes*** **$M1_{UK}$ isogenic strains nasopharyngeal carriage and shedding** Quantification of *S. pyogenes* nasal shedding as mean number of CFU recovered by daily nose pressing onto individual blood agar plates (A) and longevity of nasal shedding (B) comparing infection with isogenic $M1_{UK}$, $M1_{UKrofAKo}$ and $M1_{UKrofA3SNPsFixed (F)}$ strains (n= 5 per group). Note reduction in group size day 4 onwards. Quantification of *S. pyogenes* shedding onto settle plates on days 0-3 of infection (n=4 plates/cage) as metric of shedding into air (C).

**Figure S8**

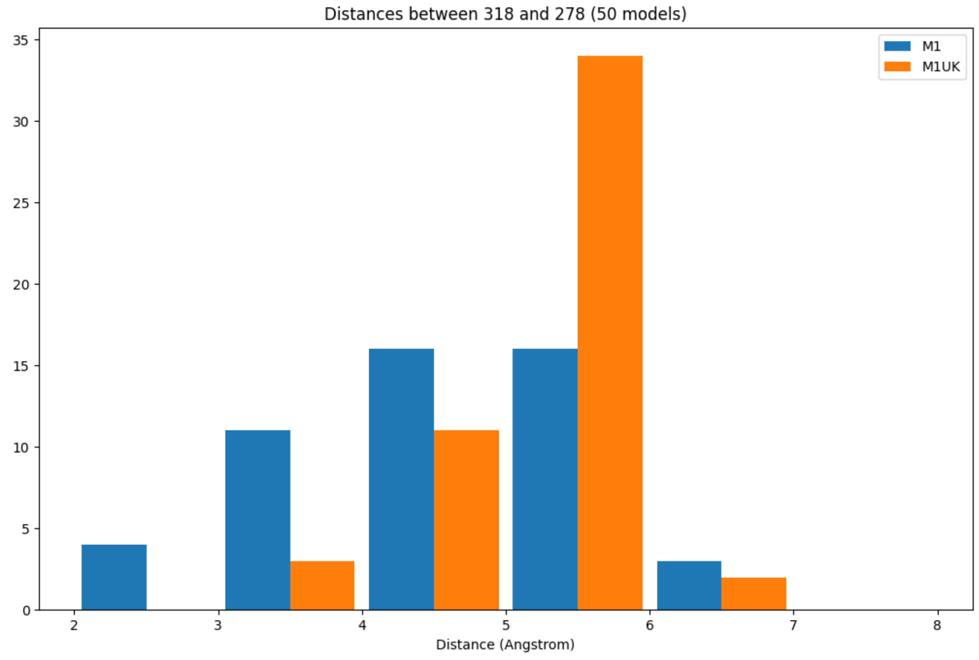


**Supplementary Figure S8.** Distribution of minimum interatomic distances between residues His-278 and Met-318 in $M1_{global}$ and $M1_{UK}$ models
